# Backup transcription factor binding sites protect human genes from mutations in the promoter

**DOI:** 10.1101/2023.01.27.525856

**Authors:** Jay C. Brown

**Affiliations:** Department of Microbiology, Immunology and Cancer Biology University of Virginia School of Medicine Charlottesville, Virginia, 22908

**Keywords:** gene expression, transcription factor, promoter, tissue specific human genes, mutation in promoter

## Abstract

This study was designed to test the idea that human gene promoters have evolved to be resistant to the effects of mutations in their primary function, the control of gene expression. It is proposed that the transcription factor/transcription factor binding site (TF/TFBS) pair having the greatest effect on control of a gene is the one with the highest abundance in the promoter. Other pairs would have the same effect on gene expression and would predominate in the event of a mutation in the most abundant pair. It is expected that the overall promoter architecture proposed here will be highly resistant to mutagenic change that would otherwise affect expression of the gene. The idea was tested beginning with a database of 42 human genes highly specific for expression in brain. For each gene, information was accumulated about its expression level and about the TFBS occupancy of the five most abundant TF/TFBS pairs. Expression level was then plotted against TFBS occupancy separately for each of the five pairs, and the plots were compared with each other. The plots were found to be similar, and the results were interpreted to indicate that the TFBS occupancy ranks evolved to yield the same effect on gene expression level with multiple ranks able to function in the event of mutation in another. A similar analysis was conducted with a database of 31 human liver specific genes, and the overall result was found to be the same. Backup TFBS occupancy ranks were interpreted to be present in both brain and liver specific genes. Finally, the TFBSs in the brain specific and liver specific gene populations were compared with each other with the goal of identifying any brain selective or liver selective TFBSs. Of the 89 TFBSs in the pooled population, 58 were found only in brain specific but not liver specific genes, 8 only in liver specific but not brain specific genes and 23 were found in both brain and liver specific genes. The results were interpreted to emphasize the large number of TFBS in brain specific but not liver specific genes.

## 1. Introduction

Investigators continue to be puzzled by the large number of transcription factor binding sites present in the promoters of human genes. Whereas in prokaryotic organisms one or a very few TF binding sites suffice for the regulatory needs of most genes, hundreds are the rule in human gene promoters. For instance, a total of 357 TFBS are annotated in DLGAP3, a human gene of average length. High TFBS numbers are also the rule in the genomes of other eukaryotes. Further, in many human genes distinct TFBS clusters are observed outside the promoter but within the coding region of a gene. Five such clusters are annotated in DLGAP3 [1,2].

Reasonable suggestions have been advanced to account for the large number of TFBS found in the genomes of eukaryotes. TFs and TFBSs have been proposed, for example, to be involved in specifying the tissue where a gene is expressed, a feature less relevant in prokaryotes [3–6]. Eukaryotic gene expression could be controlled by combinations of TFs in a way that would require an increased number of TFBSs [7–9]. Despite the availability of reasonable suggestions, however, there is currently no consensus about the role of the “extra” TF/TFBSs present in eukaryotic genomes.

The study described here was carried out to test the idea that the large number of TFBSs in eukaryotic promoters has evolved to protect the regulatory region from the effects of mutations. Since the interaction of a TFBS with its TF occurs in the promoters of many different genes, it is reasonable to expect that additional protection against mutagenic change may be required for TFBS compared to DNA regions with fewer interaction targets. The hypothesis tested in the present study proposes that the additional TFBS in eukaryotes has evolved to meet the need for the additional protection against mutation in the TF/TFBS contact.

The mechanism proposed for protection depends on the abundance of binding sites for a TF in the promoter. It is suggested that expression of a gene depends to the greatest extent on the TF that binds in the highest abundance TFBS. Lesser effects on transcription would be provided by TFBS with lower abundance in the promoter. This situation would prevail until there is a mutation causing loss of the highest abundance TF/TFBS interaction. In that case the second most abundant TFBS/TF pair would become the most abundant and gene expression would continue unchanged because the new highest abundance TFBS would dominate regulation of transcription. The proposal depends on the idea that transcription regulation depends on the identity of the highest abundance TF/TFBS pair, not on the identity of the individual TF/TFBS pair involved. A variety of different TF/TFBS pairs are proposed to be able dominate regulation of a gene if the pair is the most abundant one in the promoter.

The study described here was designed to test the idea outlined above for control of gene expression. It was reasoned that if control were indeed dominated by the most abundant TF/TFBS pair, then a recognizably similar result should be obtained if one plotted gene expression level against TFBS occupancy for the most abundant TFBS/TF pair or the second most abundant or the third and so on. If the second most abundant pair has evolved to take the place of the most abundant in the event of a mutation, then the second most abundant pair should have a similar effect on transcription as the first.

The study was performed beginning with a database of 42 human genes each with highly selective expression in the brain. Public databases were used to accumulate the expression level of each gene and the ChIP-seq signal from each of the five most abundant TFBSs. Expression and ChIP-seq results were then plotted for each TFBS abundance level with the expectation that the plots with the different abundance levels would resemble one another. Such similarity would support the view that the second most abundant rank of TFBS would have the ability to replace the first if the first were lost due to mutation.

A similar study was conducted with a database of human genes with highly selective expression in liver. Information was collected about gene expression level and TFBS occupancy for different abundance ranks as described for the brain database. Plots of expression against ChIP-seq signal were created in the same way. It was anticipated that the outcome of the liver study would clarify whether the brain results are unique to brain or if they would apply more broadly.

Finally, a study was focused on the TFBSs present in the brain database. All TFBSs in the brain population were pooled and compared to the similar pool of TFBSs in the liver group. The results were expected to identify any instances of brain-selective or liver-selective TFBSs.

## 2. Materials and Methods

### 2.1 Gene databases: gene expression and TF binding levels

Brain specific genes were accumulated for the study from a combination of locally curated genes and those from the database of Sonawane et al, [3]. Focus was on a tissue specific gene population to maximize the possibility that conclusions would apply to the level of gene expression and minimize contributions from the potential effects of TFs on tissue distribution. Tissue expression for each gene was obtained from the GTEx Portal of RNA-seq results found in the UCSC Genome Browser (version hg38 (https://genome.ucsc.edu/)). A gene was considered to be brain-specific if its expression was 10-fold or greater in brain than in the tissue with the next highest expression level. The database of liver-specific genes was curated using the same procedure described above for brain.

For each gene examined, the TF binding abundance of each TFBS was determined from the ChIP-seq values reported in the ENCODE project database of cis-Regulatory Elements (3 November 2018 version) by way of the UCSC Genome Browser. Entries were included from the promoter and from other TFBS clusters within the gene coding region. Table Browser was searched on Regulation followed by TF Clusters, and the output was focused on the “score” entry. The score was summed for each TFBS as there were several (usually ~3-10) identical TFBS in each gene. The highest scoring TFBS and its score were then recorded in the brain-specific or liver-specific database under the heading “ChIP1.” Values for the second highest TFBS were recorded under ChIP2 and so on through ChIP5. Note that the TFBS associated with the ChIP1 rank cannot be the same as that for ChIP-seq2, or ChIP-seq 3 and so on. For each gene, each ChIP-seq rank has its own TFBS.

### 2.2 Gene databases

TFBSs in brain specific and liver specific gene promoters Two TFBS populations were accumulated for the brain specific genes, (1) all TFBSs present in the database (i.e., 5 for each gene) and (2) all non-redundant TFBSs in the database. The same two populations were accumulated for the liver-specific genes.

### 2.3 Data analysis

Data were manipulated with RStudio and Excel. Graphic rendering was done with SigmaPlot v14.5.

## 3. Results

### 3.1 Experimental strategy

The analyses described here were designed to test the idea that the promoter regions of human genes have evolved to create redundant layers of TFBS groups able to substitute for one another in the event of mutagenic damage to the promoter. Relevant TFBS groups were identified according to the abundance of their occupancy by the cognate TF as determined from the results of ChIP-seq studies. For each gene, the highest abundance TFBS/TF pair was put into one pool (rank 1), the second most abundant was put in a second pool (rank 2) and so on to rank 5. Each data point therefore has its own TF and its own level of TFBS occupancy. If the information described above is to function in a redundant or backup manner to control the level of gene expression, then it is expected that each TFBS/TF rank will have the same effect on the level of gene expression. This is the expectation that was tested in the studies described below. It was assumed that a positive result would support the existence of redundant layers of regulatory control in human gene promoters.

### 3.2 Effect of TFBS occupancy rank on brain specific gene expression

The effect of TFBS occupancy rank on gene expression was examined with a population of 42 protein coding, human, brain specific genes (see Supplementary Table 1). For each occupancy rank the ChIP-seq signal was plotted against the gene expression level, and the results for the first three ranks are shown in Fig. 1. Clear similarities were observed among the three ranks in the expression/ChIP-seq signal relationship. For instance, in the lower left of the plot the RTP1, PNMA6F and FGF3 genes are found in similar locations in the three abundance ranks. Near the center of the plot, HTR5A, HCRT and TLX3 are located similarly. CACNG8, TBR1 and CREG2 locations are related in all three plots. The results are interpreted to indicate that the three ChIP-seq abundance ranks have the potential to serve in a redundant fashion to drive expression of the genes to the same level. It is suggested that the small differences observed in ChIP-seq signal for the same gene in separate ranks might produce tolerable results if one rank needed to be substituted for another.

**Fig. 1:**
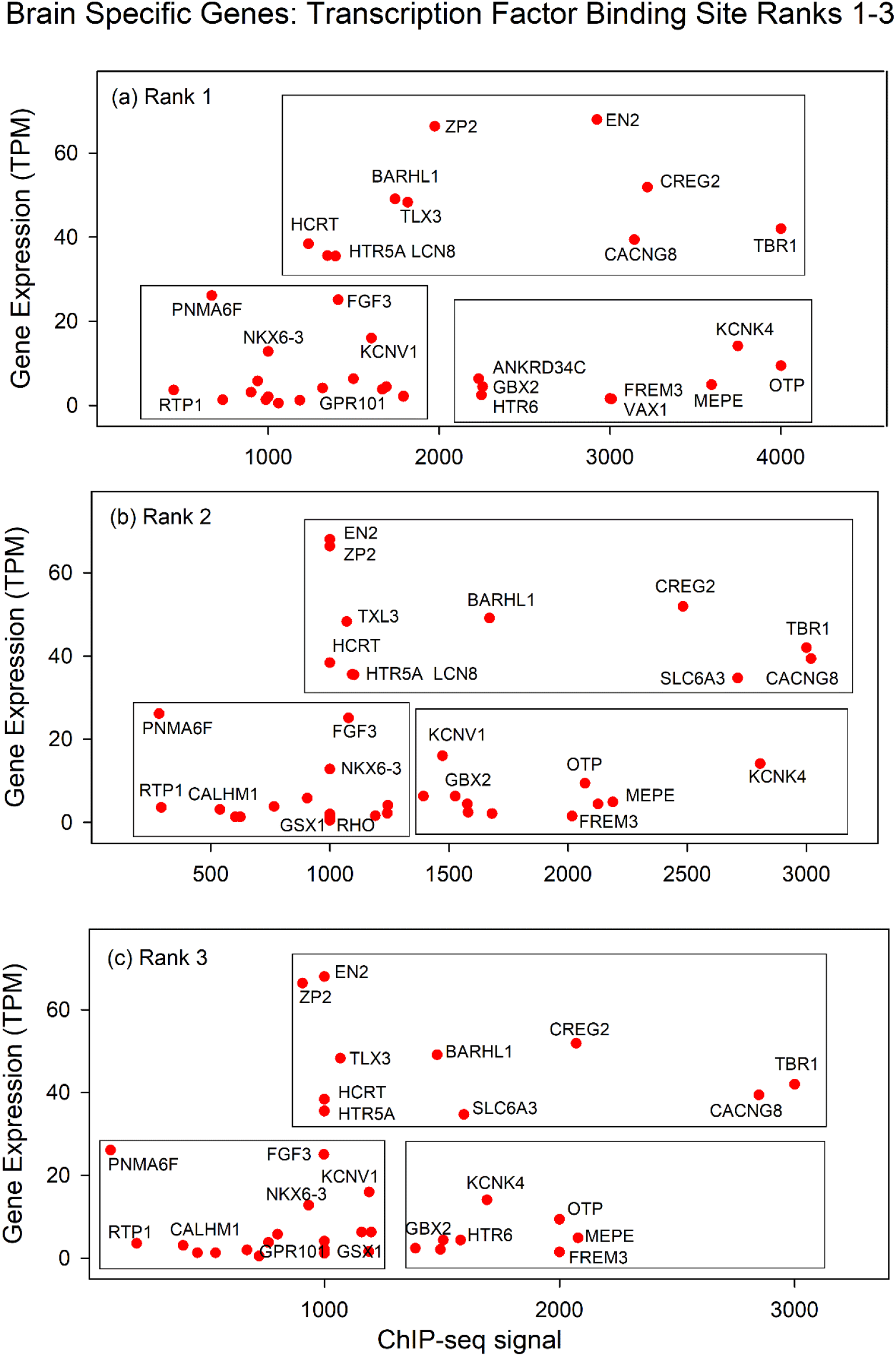
Plots of gene expression level against ChIP-seq signal for brain specific genes. Results for Ranks 1, 2, and 3 are shown in panels (a), (b) and (c), respectively. Numerical values for the data points shown can be found in Supplementary Table 1. Boxes are intended to aid in interpretation of the results. They do not contain any experimental results. Note that data points for the same gene are related in the three plots.

### 3.3 Effect of TFBS occupancy rank on liver specific gene expression

To test the generality of the results described above with brain specific genes, a similar analysis was carried out with liver specific genes. A population of 31 liver specific human genes was first identified, and each gene was assigned to a set of five ranks based on TFBS occupancy in the promoter (see Supplementary Table 2). The occupancy score, as measured by the ChIP-seq signal, was then plotted against the gene expression level.

The results showed that the same genes were found in similar locations in the three plots (see Fig. 2). CFHTR3 and UGT2B10, for instance, are located in the high expression part of each plot with CFHR3 at the lower ChIP-seq signal end. An arc of four genes with varying expression levels is found in all three plots near the low ChIP-seq signal level (see SPP2, SERPINA7, F13B and SLC22A10). All plots also have a cluster of four genes at the high end of the ChIP-seq axis (i.e., genes MBL2, INHBC, SLC22A25 and INS-IGF2). The results are interpreted to indicate that the three ChIP-seq abundance ranks can serve in a redundant or backup manner to shield expression of the gene from mutagenic change in the promoter. Further, the results show that the same backup promoter design is found in liver specific as well as brain specific human genes.

**Fig. 2:**
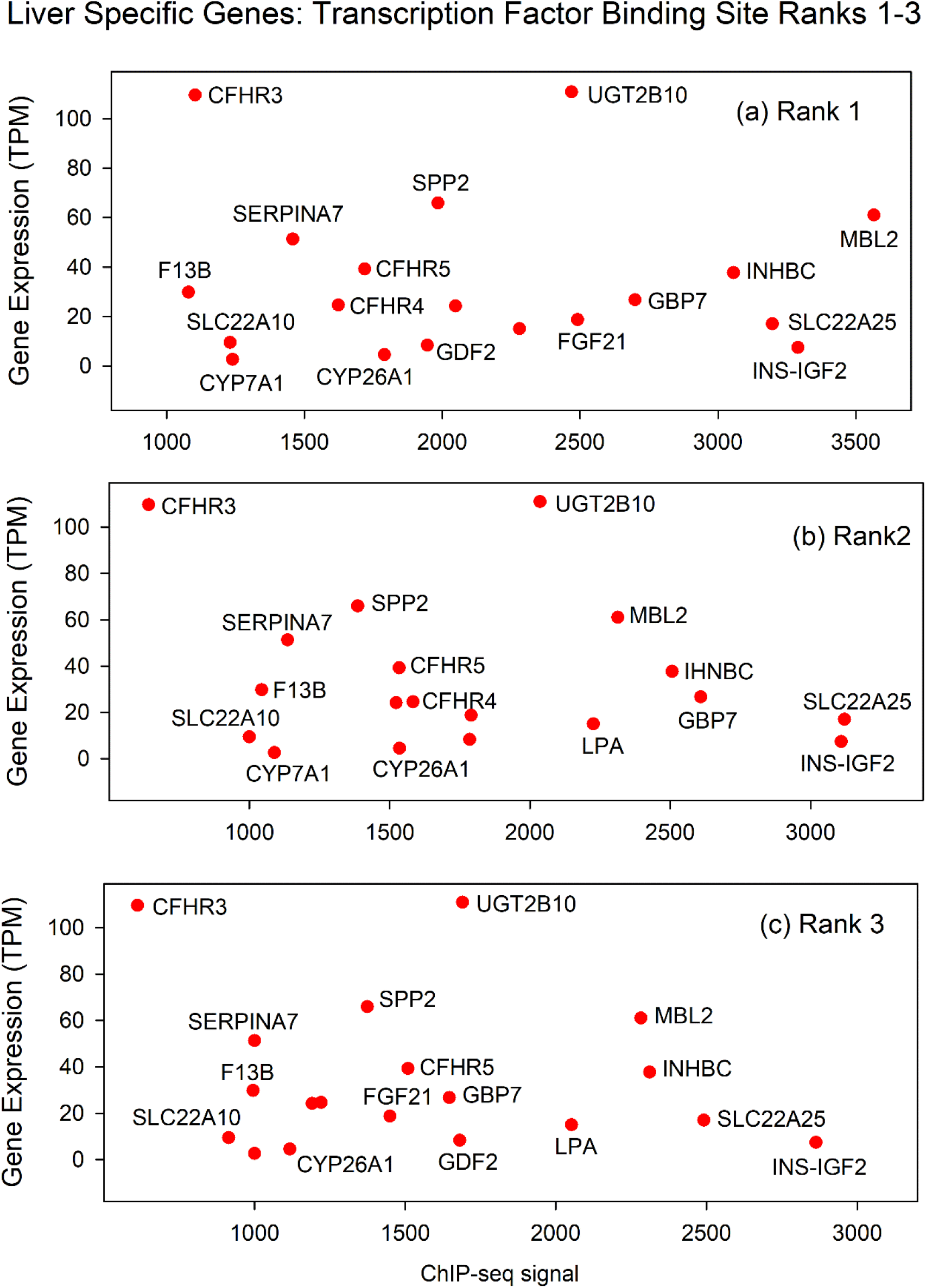
Plots of gene expression level against ChIP-seq signal for liver specific genes. Results for Ranks 1,2 and 3 are shown in panels (a), (b) and (c), respectively. Numerical values for the data points shown can be found in Supplementary Table 2. Note that the data points for the same gene are related in the three plots.

### 3.4 TFBSs in the promoters of brain and liver specific human genes: all database TFBSs

The information accumulated here about promoter content of TFBSs provides the opportunity to compare the promoters of brain specific and liver specific genes (see Supplementary Tables 1 and 2). While previous studies have demonstrated a degree of tissue selectivity to TFBSs, it is rare for such studies to involve the large populations of tissue-specific genes available here [10]. For analysis, TFBSs in the promoters of brain specific genes were assembled in two forms, (1) all the TFBSs shown in Supplementary Table 1. That is, the five most abundant TFBSs in each of the 42 brain specific genes or 210 TFBSs in all: and (2) the population of non-redundant TFBSs. Each TFBS is counted only once regardless of how many times it occurs in the overall population. The same two TFBS populations were assembled for the liver-specific genes. Finally, histograms were plotted to show the count of each TFBS in the total population (Figs. 3 and 4).

**Fig. 3:**
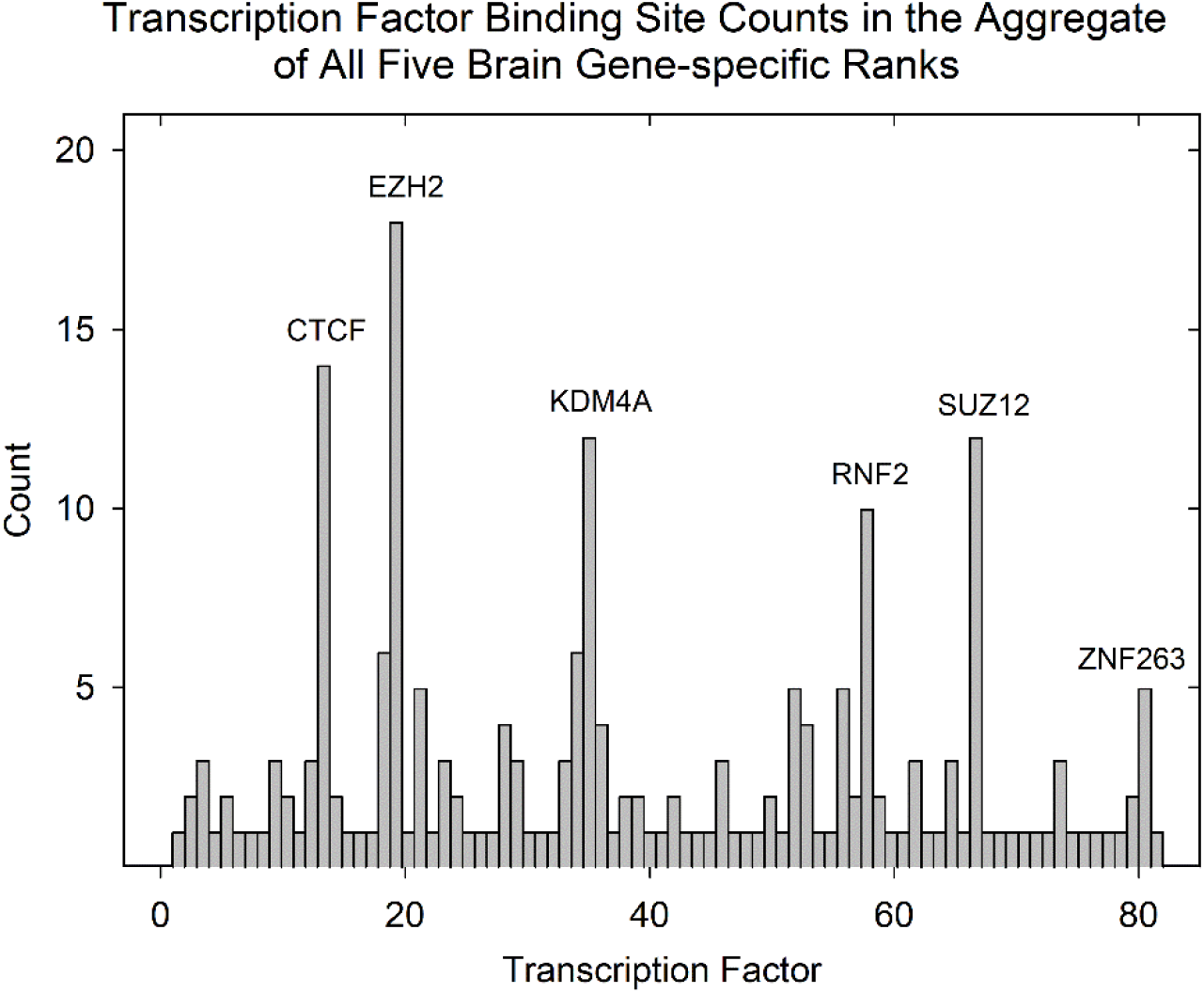
Histogram showing the count of each TFBS annotated in the brain specific gene population (see Supplementary Table 1). TFBS in all five ranks are included. TFBS not identified in the figure are noted in Supplementary Table 3. Note that 3 of the 6 TFBS with the highest abundance bind TF that are subunits of the Polycomb Repressive Complex 2 involved in histone methylation (i.e., EZH2, RNF2 and SUZ12).

**Fig 4:**
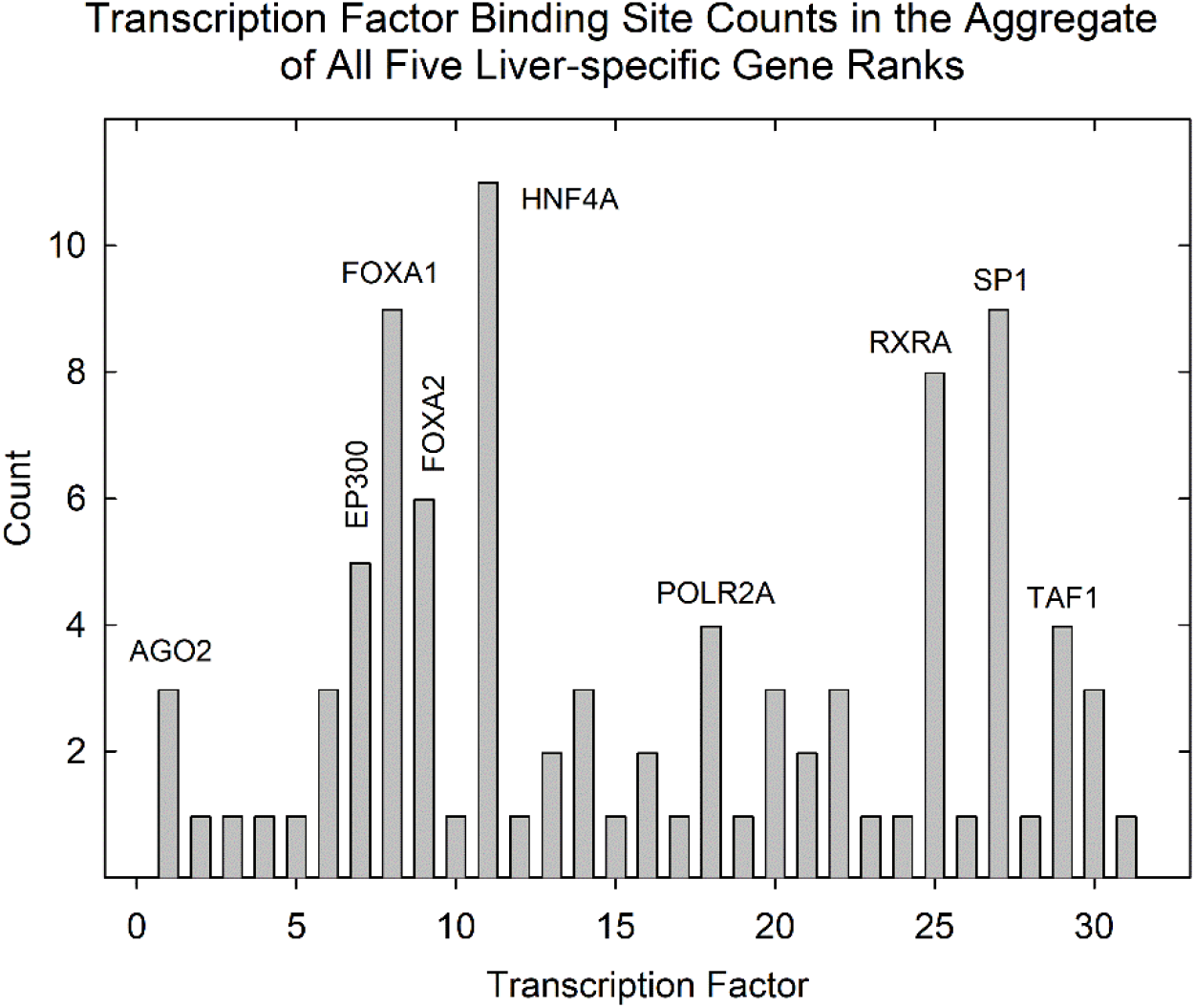
Histogram showing the count of each TFBS annotated in the liver specific gene population (see Supplementary Table 2). TFBS in all five ranks ae included. TFBS not identified in the figure are noted in Supplementary Table 3. Note the high abundance of HNF4A binding sites in the liver specific gene population.

The results for brain specific genes emphasize binding of TFs that use histone methylation to influence gene expression (Fig. 3). For instance, of the five most abundant TFBSs three (EZH2, SUZ12 and RNF2) are components of polycomb repressive complex 2 (PRC2), a complex that uses histone methylation to attenuate gene expression [11, 12]. A fourth, KDM4A, also uses histone methylation to regulate transcription [13]. The result points to an important role for histone methylation in the regulation of brain specific genes.

No similar emphasis on histone methylation was observed with liver-specific genes (Fig. 4). Of the seven most abundant TFBSs, three bind TFs that attract transcription machinery by binding to the promoter region (i.e., FOXA1, FOXA2 and HNF4A) [14–16]. Three of the four remaining TFBSs affect transcription of a wide variety of genes. RXRA mediates the effects of retinoids while SP1 and YY1 can activate or repress gene expression depending on other factors [17–19]. Comparing the brain and liver specific gene populations suggests the dominant mechanisms of gene regulation are distinct in the two tissues.

### 3.5 TFBSs in the promoters of brain and liver specific human genes: all unique database TFBSs

Non-redundant TFBS in the brain and liver specific databases were accumulated with the expectation that they might identify TFBSs found in brain-specific genes, but not in liver specific ones. TFBS found in liver specific, but not brain specific genes would also be identified. Scans of Supplementary Tables 1 and 2 revealed the presence of 81 non-redundant TFBS among the 210 total brain-specific genes and 31 among the 155 liver-specific genes (Table 1). Percentages were 39% in the case of brain specific genes and 20% for liver. Among the 81 non-redundant brain TFBS, 58 (72%) were found in the brain population only and 23 (28%) in both brain and liver populations. In the 31 non-redundant TFBS in liver, 8 (26%) were found in liver, but not brain and 23 (74%) in both liver and brain.

**Table 1:**
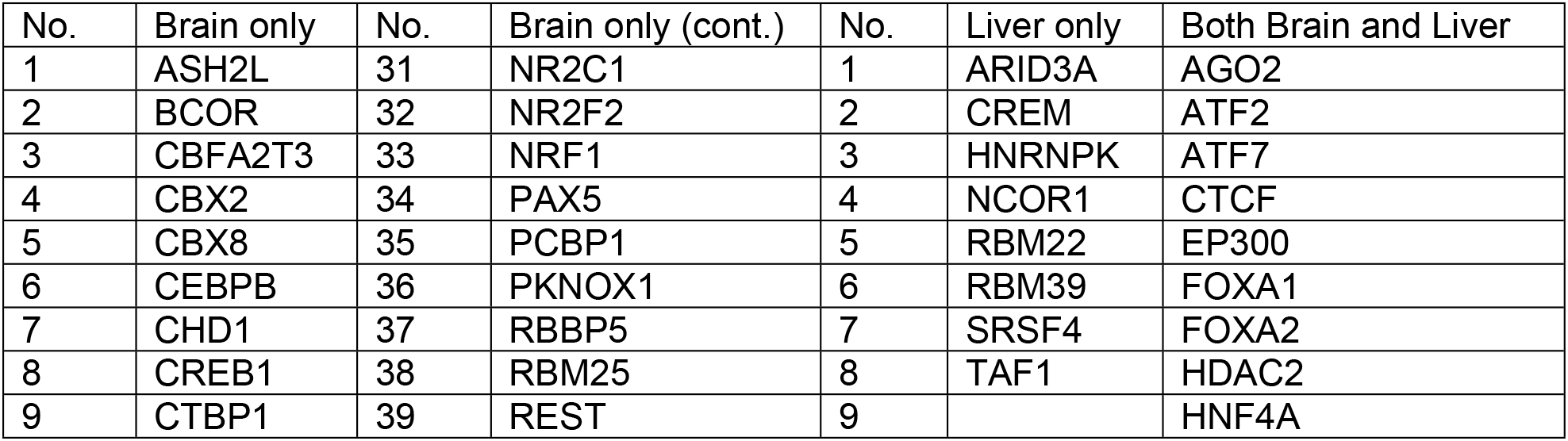
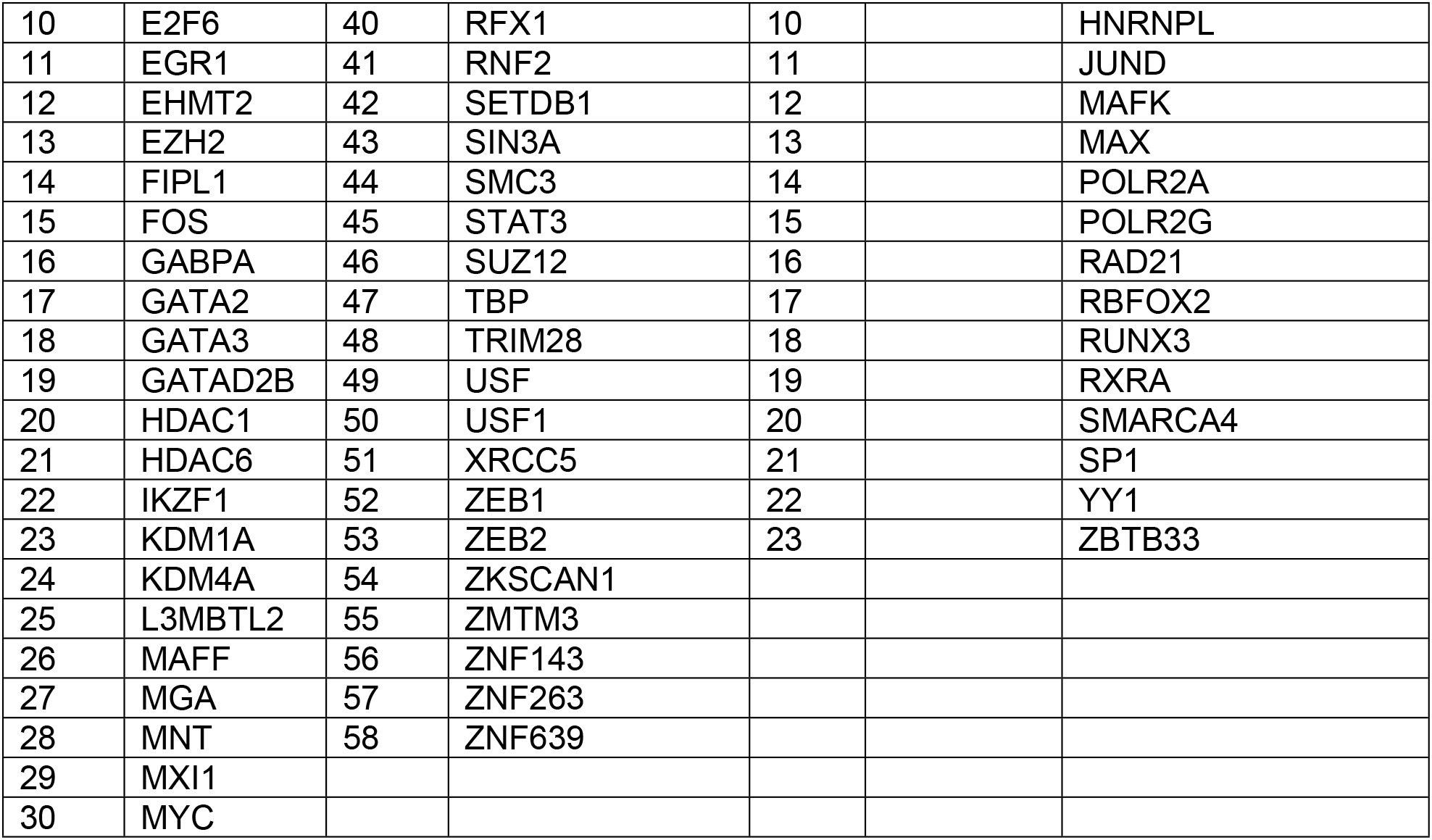
Transcription Factor Binding Sites Present in the Promoters of Database Brain-specific and Liver-specific Genes (all Five Abundance Ranks)

Two conclusions stand out from analysis of the non-redundant TFBS populations described above. First is the high proportion of non-redundant TFBS among the brain specific genes (58 in the 81 total non-redundant TFBS population). For liver, the comparable figures are 8 of 31 non-redundant TFBS. The high proportion among brain genes may result from a greater need for gene regulatory activity in the brain due perhaps to a greater number of functionally distinct regions in the brain or to a more diverse developmental program.

Second is the absence of any evidence for a tissue specific TFBS. All the TFBS found in the brain-specific gene population, for instance, are those that can be readily identified in the promoters of non-brain genes. The same is true for the TFBS identified here in liver specific, but not brain specific genes (Table 1). All can be found abundantly in other human genes. The overall finding indicates that the role of TF in influencing the tissue where a gene is expressed must involve multiple TFs or TFs in combination with other regulatory mechanisms.

## 4. Discussion

### 4.1 Backup organization to allow promoter function to co-exist with mutagenic damage

The backup system of promoter organization suggested by the results described here is distinct from other methods humans use to deal with mutagenic change to the genome. The suggested system does not result in prevention of mutations or correction of them once they have occurred. Rather, the system describes a method promoters use to continue to function normally despite the existence of mutations in their own sequences. The results indicate the existence of backup or redundant systems of TFBS that are each able to provide gene regulatory function in the event of mutagenic damage to another. An individual promoter is thereby able to co-exist with a diverse background of mutagenic events using methods that complement other mechanisms that prevent or correct mutations.

The backup system can be understood as related to other genome-wide features that are able to allow the individual to accommodate potentially disastrous mutagenic events. An example is the way genes encoding proteins that are parts of a single structure or function are distributed widely in the genome rather than being together in a cluster. The 31 genes encoding distinct protein components of the human nuclear pore complex are an example. The genes are spread among 17 different chromosomes increasing the likelihood that damage to any one gene might be able to be accommodated by the overall structure [20, 21]. This arrangement prevents a mutation in a single chromosome from affecting more than one or a few nuclear pore genes. Other distributed genomic elements that enable humans to co-exist in a mutagenic environment include gene enhancers, homologous genes and exons distributed within the same gene [22–25].

### 4.2 High TFBS content of eukaryotic promoters

The system of backup TFBSs proposed here is consistent with and supports the observation that the promoters of human genes and those of other eukaryotic organisms contain many more TFBSs than those of prokaryotes. The ChIP-seq signal from the most abundant TFBS is in most cases a sum of several individual TFBSs in the promoter. The same is true of the second most abundant TFBS, the third and so on. It is easy to imagine therefore that the number of individual TFBSs in the aggregate of all abundance classes (i.e. ranks) might reach the high number observed in an entire gene promoter region.

### 4.3 Tissue specific genes

This study was focused on tissue specific genes with the idea that the results might clarify the way tissue specificity is encoded in promoters. The results confirmed a previously noted selectivity of HNF4A for liver genes [26]. A binding site for HNF4A was observed to be among the five most abundant in 19 of the 31 liver specific genes examined here (see Supplementary Table 2). Evidence for other tissue specific TFBSs was less convincing. For instance, 21 different TFBS were found among the 42 highest abundance TFBS in the brain specific genes examined here (see Supplementary Table 1). Overall, the results provide little support for the view that there is a single TF that uniquely determines the tissue location of a brain or liver specific human gene.

### 4.4 Role of TFBS occupancy in the level of gene expression

The results described here provide a test of the idea that the level of TF binding to its TFBS may influence the level of a gene’s expression, at least in some cases. Support for the idea would be obtained if the level of TFBS occupancy as measured by the ChIP-seq signal were found to be correlated with the level of gene transcription. Such correlations can be found in both the brain and liver gene populations examined here (see Figs. 1 and 2).

Among the brain genes an inverse correlation is observed among the genes ZP2, BARHL1, CREG2, CACNG8 and TBR1 in all three abundance ranks plotted. In the liver gene population, a positive correlation is observed relating genes F13B, SERPIN7, SPP2 and UGT2B10 in all three ranks shown. The results suggest that expression of genes in the brain group is repressed by TFBS occupancy in all three TF ranks while expression is activated in the liver gene group. In both cases the results support the view that interaction of TF and TFBS can have an influence on transcription level in the genes involved.

## Supplementary Tables

**Supplementary Table 1:** AH human brain-specific genes used in this study (42 genes)

| gene | tfbs1<sup>a</sup> | ChIP1<sup>b</sup> | tfbs2 | ChIP2 | tfbs3 | ChIP3 | tfbs4 | ChIP4 | tfbs5 | ChIP5 | exp1<sup>c</sup> |
| --- | --- | --- | --- | --- | --- | --- | --- | --- | --- | --- | --- |
| ANKRD34C | REST | 2231 | CTCF | 1526 | EHMT2 | 1158 | EZH2 | 1000 | HDAC2 | 1000 | 6.3 |
| ANKRD63 | HNRNPL | 1691 | HDAC2 | 1577 | CTBP1 | 1505 | KDM4A | 1478 | L3MBTL2 | 1327 | 4.4 |
| BARHL1 | RNF2 | 1743 | SUZ12 | 1670 | REST | 1479 | EZH2 | 1000 | HDAC1 | 1000 | 49.1 |
| BHLHA9 | EZH2 | 735 | HDAC6 | 604 | KDM4A | 460 | GABPA | 460 | YY1 | 310 | 1.3 |
| BTBD17 | ZEB1 | 1499 | ZNF143 | 1393 | CTCF | 1199 | GABPA | 1000 | NR2C1 | 1000 | 6.3 |
| C4orf50 | CTCF | 16230 | RAD21 | 11686 | KDM1A | 7288 | ZNF143 | 6682 | MXI1 | 6400 | 9.7 |
| CACNG8 | SIN3A | 3143 | SP1 | 3019 | JUND | 2848 | POLR2A | 2755 | KDM4A | 2372 | 39.4 |
| CALHM1 | USF | 900 | GATAD2B | 539 | ZNF263 | 399 | MAX | 376 | ZBTB33 | 340 | 3.1 |
| CREG2 | FOS | 3219 | CTCF | 2482 | MAFK | 2071 | MAFF | 1734 | RAD21 | 1509 | 51.9 |
| EN2 | RNF2 | 2924 | EZH2 | 1000 | SUZ12 | 1000 | ZNF263 | 1000 | L3MBTL2 | 614 | 68 |
| FGF3 | EZH2 | 1410 | SUZ12 | 1079 | KDM4A | 998 | GATA2 | 795 | RXRA | 775 | 25.1 |
| FREM3 | EP300 | 3009 | CTCF | 2017 | RUNX3 | 2000 | FOS | 1688 | NR2F2 | 1586 | 1.5 |
| GBX2 | RNF2 | 2248 | POLR2A | 1580 | E2F6 | 1387 | EGR1 | 1330 | USF1 | 1000 | 2.4 |
| GPR101 | KDM4A | 1319 | EZH2 | 1244 | NRF1 | 1000 | SUZ12 | 865 | L3MBTL2 | 647 | 4.1 |
| GRIN2B | CTCF | 11057 | SP1 | 11020 | EP300 | 10847 | JUND | 9741 | RAD21 | 8550 | 6.9 |
| GRM4 | CTCF | 12716 | RBM25 | 6865 | KDM1A | 6510 | ZMYM3 | 6404 | RAD21 | 6323 | 286.8 |
| GSX1 | REST | 1185 | EZH2 | 1000 | RNF2 | 1000 | SIN3A | 1000 | SUZ12 | 1000 | 1.2 |
| HCRT | IKZF1 | 1236 | L3MBTL2 | 1000 | MAX | 1000 | MGA | 1000 | MYC | 1000 | 38.4 |
| HTR5A | REST | 1394 | MXI1 | 1101 | TRIM28 | 1000 | XRCC5 | 1000 | ZBTB33 | 1000 | 35.5 |
| HTR6 | KDM1A | 2254 | REST | 2126 | SIN3A | 1579 | EHMT2 | 1177 | NRF1 | 1051 | 4.4 |
| KCNK4 | EZH2 | 3748 | SUZ12 | 2806 | RNF2 | 1691 | KDM4A | 1167 | ZKSCAN1 | 1000 | 14.1 |
| KCNV1 | KDM4A | 1604 | EZH2 | 1473 | HDAC2 | 1190 | CTCF | 1147 | SUZ12 | 944 | 16 |
| LCN8 | ZMTM3 | 1347 | CTCF | 1094 | BCOR | 1000 | KDM1A | 1000 | SETDB1 | 1000 | 35.6 |
| MEPE | FOS | 3594 | CTCF | 2188 | STAT3 | 2079 | RAD21 | 1870 | SMC3 | 1210 | 4.9 |
| MINDY4B | EP300 | 1791 | FOS | 1681 | CEBPB | 1493 | GATA3 | 1468 | CBX8 | 1436 | 2.1 |
| NKX6-3 | KDM4A | 1000 | RBBP5 | 1000 | ASH2L | 933 | EZH2 | 856 | HDAC6 | 721 | 12.8 |
| OTP | RNF2 | 4000 | PAX5 | 2071 | SUZ12 | 2000 | RBBP5 | 1894 | RUNX3 | 1823 | 9.4 |
| PABPN1L | AGO2 | 1668 | PCBP1 | 766 | FIPL1 | 762 | RBFOX2 | 657 | POLR2G | 541 | 3.8 |
| PNMA6F | KDM4A | 670 | ZEB2 | 284 | RFX1 | 91 | PKNOX1 | 63 | CBFA2T3 | 28 | 26.1 |
| POU6F2 | SMARCA4 | 16814 | EP300 | 11208 | CTCF | 10901 | MAFK | 7561 | EZH2 | 6618 | 1.9 |
| PTH2 | ZBTB33 | 1791 | KDM1A | 1241 | CTBP1 | 1000 | CTCF | 1000 | EZH2 | 1000 | 2.2 |
| RHO | SP1 | 1060 | CEBPB | 1000 | ASH2L | 722 | RBBP5 | 710 | CTBP1 | 653 | 0.5 |
| RTP1 | NRF1 | 448 | EP300 | 293 | ZNF263 | 202 | ZNF639 | 172 | RFX1 | 169 | 3.6 |
| SLC35D3 | EZH2 | 939 | HDAC6 | 905 | KDM4A | 801 | SUZ12 | 627 | E2F6 | 492 | 5.8 |
| SLC6A3 | EZH2 | 7848 | MNT | 2712 | CTCF | 1593 | HDAC2 | 1573 | ATF2 | 1472 | 34.7 |
| SSTR4 | KDM4A | 987 | EZH2 | 625 | ZNF263 | 537 | SUZ12 | 457 | GABPA | 317 | 1.3 |
| STH | ATF2 | 1000 | ATF7 | 1000 | JUND | 671 | CREB1 | 633 | BCOR | 495 | 2 |
| TBR1 | EZH2 | 4000 | RNF2 | 3000 | SUZ12 | 3000 | CBX2 | 2542 | CHD1 | 2105 | 42 |
| TLX3 | RNF2 | 1816 | RBBP5 | 1071 | TBP | 1068 | EZH2 | 1000 | ZNF263 | 1000 | 48.3 |
| TMEM132D | FOS | 14312 | EP300 | 12091 | CTCF | 11202 | ATF2 | 8464 | KDM1A | 8448 | 9 |
| VAX1 | RNF2 | 3000 | CHD1 | 1191 | KDM4A | 1188 | EZH2 | 1000 | SUZ12 | 1000 | 1.6 |
| ZP2 | CEBPB | 1974 | RNF2 | 1000 | HNF4A | 907 | GATA2 | 748 | FOXA2 | 669 | 66.4 |

<sup>a</sup> Most abundant transcription factor binding site (i.e., rank 1 transcription factor binding site).
<sup>b</sup> ChIP-seq signal for rank 1 transcription factor binding site.
<sup>c</sup> Gene expression in TPM from GTEx.

**Supplementary Table 2:** All human liver-specific genes used in this study (31 genes)

| gene | tfbs1<sup>a</sup> | ChIP1<sup>b</sup> | tfbs2 | ChIP2 | tfbs3 | ChIP3 | tfbs4 | ChIP4 | tfbs5 | ChIP5 | exp<sup>c</sup> |
| --- | --- | --- | --- | --- | --- | --- | --- | --- | --- | --- | --- |
| AFP | POLR2A | 9187 | RBM22 | 8670 | RBFOX2 | 2310 | FOXA2 | 1858 | FOXA1 | 1819 | 1.3 |
| AGXT | HNRNPL | 4336 | RXRA | 4203 | HNF4A | 4200 | MAX | 3624 | SP1 | 3575 | 1731.5 |
| AHSG | RBM22 | 5881 | POLR2A | 5374 | AGO2 | 4172 | RBM39 | 3775 | SRSF4 | 3394 | 1346.5 |
| C8A | HNF4A | 8689 | EP300 | 7435 | JUND | 5913 | MAX | 5901 | FOXA1 | 5489 | 274 |
| C8B | HNF4A | 5517 | SP1 | 4501 | FOXA2 | 2983 | RXRA | 2968 | FOXA1 | 2579 | 295.2 |
| C9 | HNF4A | 5403 | RXRA | 4667 | SP1 | 4659 | TAF1 | 4198 | YY1 | 3673 | 404.9 |
| CA5A | AGO2 | 8657 | RBM22 | 6267 | CTCF | 5422 | RAD21 | 3852 | POLR2A | 3659 | 6.4 |
| CFHR2 | CBX5 | 597 | RELB | 472 | RUNX3 | 451 | BATF | 444 | NFIC | 368 | 298.7 |
| CFHR3 | TAF1 | 1104 | RXRA | 641 | HNF4A | 612 | YY1 | 335 | POLR2A | 321 | 109.6 |
| CFHR4 | CTCF | 1623 | RAD21 | 1583 | YY1 | 1221 | HNF4A | 1199 | FOXA1 | 1140 | 24.6 |
| CFHR5 | YY1 | 1719 | FOXA2 | 1534 | RXRA | 1509 | HNF4A | 1318 | JUND | 1162 | 39.2 |
| CYP26A1 | HDAC2 | 1790 | ZBTB33 | 1535 | RXRA | 1117 | NCOR1 | 1000 | TAL1 | 1000 | 4.5 |
| CYP7A1 | JUND | 1240 | FOXA2 | 1089 | ARID3A | 1000 | FOXA1 | 1000 | HNF4A | 1000 | 2.6 |
| F13B | RUNX | 1080 | FOXA1 | 1044 | RAD21 | 995 | RAD21 | 893 | TAF1 | 789 | 29.8 |
| F2 | RBM22 | 7303 | AGO2 | 6184 | RBFOX2 | 5977 | POLR2A | 5356 | POLR2G | 4776 | 599.7 |
| F9 | HNF4A | 2685 | SP1 | 2607 | RXRA | 2378 | FOXA2 | 2294 | FOXA1 | 1759 | 235.2 |
| FGF21 | RXRA | 2491 | SP1 | 1790 | HNF4A | 1449 | POLR2A | 1413 | EP300 | 1177 | 18.7 |
| GBP7 | YY1 | 2699 | HNF4A | 2609 | SP1 | 1646 | MAFK | 1355 | FOXA1 | 1285 | 26.7 |
| GDF2 | RNF2 | 1946 | UBTF | 1785 | MNT | 1681 | MAX | 1449 | L3MBTL2 | 1431 | 8.3 |
| INHBC | HNF4A | 3056 | MAX | 2507 | EGR1 | 2311 | RXRA | 2128 | NR2F2 | 2111 | 37.7 |
| INS-IGF2 | CTCF | 3290 | RAD21 | 3109 | SMC3 | 2863 | MAX | 1653 | RXRA | 1377 | 7.4 |
| LPA | SMARCA | 2280 | CTCF | 2226 | ATF2 | 2052 | YY1 | 1687 | ATF7 | 1635 | 15 |
| MBL2 | RXRA | 3565 | FOXA1 | 2313 | TAF1 | 2282 | FOXA2 | 2247 | EP300 | 2000 | 61 |
| RTP3 | RXRA | 2048 | CEBPB | 1523 | HNF4G | 1191 | HNF4A | 1187 | SP1 | 968 | 24.2 |
| SERPINA7 | RBFOX2 | 1458 | RAD21 | 1136 | CTCF | 1000 | HNF4A | 1000 | MAFF | 1000 | 51.3 |
| SLC17A2 | HNF4A | 4963 | FOXA1 | 3322 | EP300 | 2969 | FOXA2 | 2960 | SP1 | 2607 | 34.5 |
| SLC22A10 | FOXA1 | 1231 | HNF4A | 1000 | FOXA2 | 914 | FOSL2 | 868 | ARID3A | 846 | 9.4 |
| SLC22A25 | FOXA2 | 3197 | SP1 | 3121 | HNF4A | 2491 | EP300 | 2221 | FOS | 2172 | 17 |
| SPP2 | HNF4A | 1984 | RXRA | 1386 | JUND | 1374 | HNRNPK | 1259 | SP1 | 1196 | 65.9 |
| UGT2B10 | FOXA2 | 2469 | RP300 | 2036 | FOXA1 | 1690 | CREM | 1383 | RBFOX2 | 1263 | 110.9 |
| UROC1 | SP1 | 4222 | CTCF | 3664 | RAD21 | 2591 | HNF4A | 2229 | HNRNPL | 2065 | 75.6 |

<sup>a</sup> Most abundant transcription factor binding site (i.e., rank 1 transcription factor binding site).
<sup>b</sup> ChIP-seq signal for rank 1 transcription factor binding site.
<sup>c</sup> Gene expression in TPM from GTEx.

**Supplementary Table 3:** Key to Transcription Factor Binding Site Numbering in Figurers 3 and 4.

| No. | Brain (Fig. 3) | Liver (Fig. 4) |
| --- | --- | --- |
| 1 | AGO2 | AGO2 |
| 2 | ASH2L | ARID3A |
| 3 | ATF2 | ATF2 |
| 4 | ATF7 | ATF7 |
| 5 | BCOR | CREM |
| 6 | CBFA2T3 | CTCF |
| 7 | CBX2 | EP300 |
| 8 | CBX8 | FOXA1 |
| 9 | CEBPB | FOXA2 |
| 10 | CHD1 | HDAC2 |
| 11 | CREB1 | HNF4A |
| 12 | CTBP1 | HNRNPK |
| 13 | CTCF | HNRNPL |
| 14 | E2F6 | JUND |
| 15 | EGR1 | MAFK |
| 16 | EHMT2 | MAX |
| 17 | EP300 | NCOR1 |
| 18 | EZH2 | POLR2A |
| 19 | FIPL1 | POLR2G |
| 20 | FOS | RAD21 |
| 21 | FOXA1 | RBFOX2 |
| 22 | FOXA2 | RBM22 |
| 23 | GABPA | RBM39 |
| 24 | GATA2 | RUNX3 |
| 25 | GATA3 | RXRA |
| 26 | GATAD2B | SMARCA4 |
| 27 | HDAC1 | SP1 |
| 28 | HDAC2 | SRSF4 |
| 29 | HDAC6 | TAF1 |
| 30 | HNF4A | YY1 |
| 31 | HNRNPL | ZBTB33 |
| 32 | IKZF1 |  |
| 33 | JUND |  |
| 34 | KDM1A |  |
| 35 | KDM4A |  |
| 36 | L3MBTL2 |  |
| 37 | MAFF |  |
| 38 | MAFK |  |
| 39 | MAX |  |
| 40 | MGA |  |
| 41 | MNT |  |
| 42 | MXI1 |  |
| 43 | MYC |  |
| 44 | NR2C1 |  |
| 45 | NR2F2 |  |
| 46 | NRF1 |  |
| 47 | PAX5 |  |
| 48 | PCBP1 |  |
| 49 | PKNOX1 |  |
| 50 | POLR2A |  |
| 51 | POLR2G |  |
| 52 | RAD21 |  |
| 53 | RBBP5 |  |
| 54 | RBFOX2 |  |
| 55 | RBM25 |  |
| 56 | REST |  |
| 57 | RFX1 |  |
| 58 | RNF2 |  |
| 59 | RUNX3 |  |
| 60 | RXRA |  |
| 61 | SETDB1 |  |
| 62 | SIN3A |  |
| 63 | SMARCA4 |  |
| 64 | SMC3 |  |
| 65 | SP1 |  |
| 66 | STAT3 |  |
| 67 | SUZ12 |  |
| 68 | TBP |  |
| 69 | TRIM28 |  |
| 70 | USF |  |
| 71 | USF1 |  |
| 72 | XRCC5 |  |
| 73 | YY1 |  |
| 74 | ZBTB33 |  |
| 75 | ZEB1 |  |
| 76 | ZEB2 |  |
| 77 | ZKSCAN1 |  |
| 78 | ZMTM3 |  |
| 79 | ZNF143 |  |
| 80 | ZNF263 |  |
| 81 | ZNF639 |  |

